# Phages Produce Persisters

**DOI:** 10.1101/2023.10.17.562728

**Authors:** Laura Fernández-García, Joy Kirigo, Daniel Huelgas-Méndez, María Tomás, Rodolfo García-Contreras, Thomas K. Wood

## Abstract

Bacteria primarily encounter stress, and, arguably, their greatest threats are phages. It is often assumed that those bacteria that escape phage attack have mutated; however, another possibility is that a subpopulation forms the dormant persister state, in a manner similar to that demonstrated for bacterial cells undergoing nutritive, oxidative, and antibiotic stress. Persister cells do not undergo mutation and survive lethal conditions by ceasing growth transiently. Slower growth and dormancy play a key physiological role as they allow host phage defense systems more time to clear the phage infection. Here we investigated how bacteria survive lytic phage infection by isolating surviving cells from the plaques of T2, T4, and lambda (cI mutant) virulent phages. We found that bacteria in plaques can escape phage attack both by mutation (i.e., become resistant) and without mutation (i.e., become persistent). Specifically, whereas T4-resistant and lambda-resistant bacteria with over a 100,000-fold less sensitivity were isolated from plaques with obvious genetic mutations (e.g., causing mucoidy), cells were also found after T2 infection that undergo no significant mutation, retain wild-type phage sensitivity, and survive lethal doses of antibiotics. Corroborating this, adding T2 phage to persister cells resulted in 137,000-fold more survival compared to that of addition to exponentially-growing cells. Phage treatments with *Klebsiella pneumonia* and *Pseudomonas aeruginosa* also generated persister cells. Hence, along with resistant strains, bacteria also form persister cells during phage infection.

## INTRODUCTION

Given the lack of novel antibiotics and the increasing mortality due to bacterial infection (∼5M deaths/yr associated with antimicrobial resistance)^16^, interest is surging in the use of phages to combat infections^26^. However, bacteria can mutate rapidly to undermine some phage therapies^14, 16^.

To date, most authors have either assumed or found that cells that survive phage attack are resistant; i.e., they assume that the host survived due to genetic change. For example, a recent guide for studying plaques of lytic phages indicates only phage-resistant bacteria are possible inside plaques (rather than transiently insensitive bacteria)^1^, and early literature reported only the formation of phage-resistant mutants of *Pseudomonas aeruginosa* spp. in the clear and confluent lysis zone^22^. In addition, neonatal meningitis *Escherichia coli* colonies resistant to lytic phage EC200^PP^ were assumed resistant, and a clearly mutant derivative (due to its rough colony morphology) was studied further and found to be less virulent^23^. Furthermore, all microcolonies in plaques were assumed to be formed by phage-resistant *Klebsiella pneumoniae* in a report for monitoring plaque growth with a wide-filed lensless imaging device^21^. Hence, the formation of persister cells as a result of phage infection has generally not been considered. A related phenotype has been reported T4 infection of stationary-phase *E. coli* cells termed ‘hibernating’; however, these hibernators were not investigated for persistence^4^, and T4 resumed lytic growth in these hibernating cells once nutrients were provided^4^.

Persister cells transiently survive myriad forms of stress based their cellular inactivity^13^. The secondary messengers guanosine pentaphosphate and guanosine tetraphosphate signal the external stress and lead to ribosome dimerization, which ceases translation in persister cells^25^. Single-cell microscopy has demonstrated persister cells resuscitate based on the re-activation of these dimerized ribosomes^11, 24, 29.^ Becoming persistent (i.e., dormant) is advantageous for bacteria to combat phages since slow growth/dormancy (i) increases time phage-defense systems to function, (ii) slows production of phage-dependent proteins and nucleic acids, (iii) increases time for spacer accumulation for CRISPR-Cas, (iv) enhances genetic diversity, and (v) reduces bacteria-phage coevolution^8^. For example, activation of the MqsR/MqsA/MqsC tripartite toxin/antitoxin system in *E. coli* by phage T2 attack results in cells entering the persister state, which enables the EcoK McrBC restriction modification system to eliminate T2 phage^6^. Moreover, expression of GTPase RsgA allows *E. coli* phage to withstand better T4 phage infection, likely as a result of persistence as a result of inactivating ribosomes^7^. In a similar fashion, the *Listeria* spp. type VI CRISPR-Cas system induces dormancy, which allows, restriction/modification systems to eliminate phage^28^. However, whether bacteria escape phage infection through persistence in general is not well studied.

Since transient resistance to phages could also undermine phage therapy by allowing pathogens to escape phage killing, we explored here whether the *E. coli* cells found inside plaques formed by T2, T4, and lambda *cI* survive by mutating or by becoming persistent. We discovered that 0.01% of the cells in suspension survive T2 infection as persister cells, rather than undergo mutation.

## RESULTS

### Kill curves with T2 phage and ampicillin

We reasoned that if persister cells form and survive during phage attack, there would be a sub-population of cells that survive in an analogous fashion to those that survive as persister cells during antibiotic killing^13^ and starvation^10^; a scheme showing the experimental plan for a series of experiments to explore this hypothesis is shown in **Fig. 1a**. We found addition of ampicillin at 10x the lethal dose (10 MIC) led to 0.1% cell survival and a clear plateau was reached in 2 h; hence, persister cells were clearly seen with ampicillin (**Fig. 1b**). Similarly, the addition of T2 phage at multiplicity of infection (MOI) of 0.1 caused a precipitous drop in cell density that plateaued with 0.001% cell survival (**Fig. 1b**); hence, since all cells did not die, persister cells are likely present. However, unlike the antibiotic-treated cells, those cells with phage addition began to recover and grow after 1 h (specific growth rate of 1.2 ± 0.1 h^-1^), which indicates resistant cells are present along with the persister cells.

**Fig. 1.**
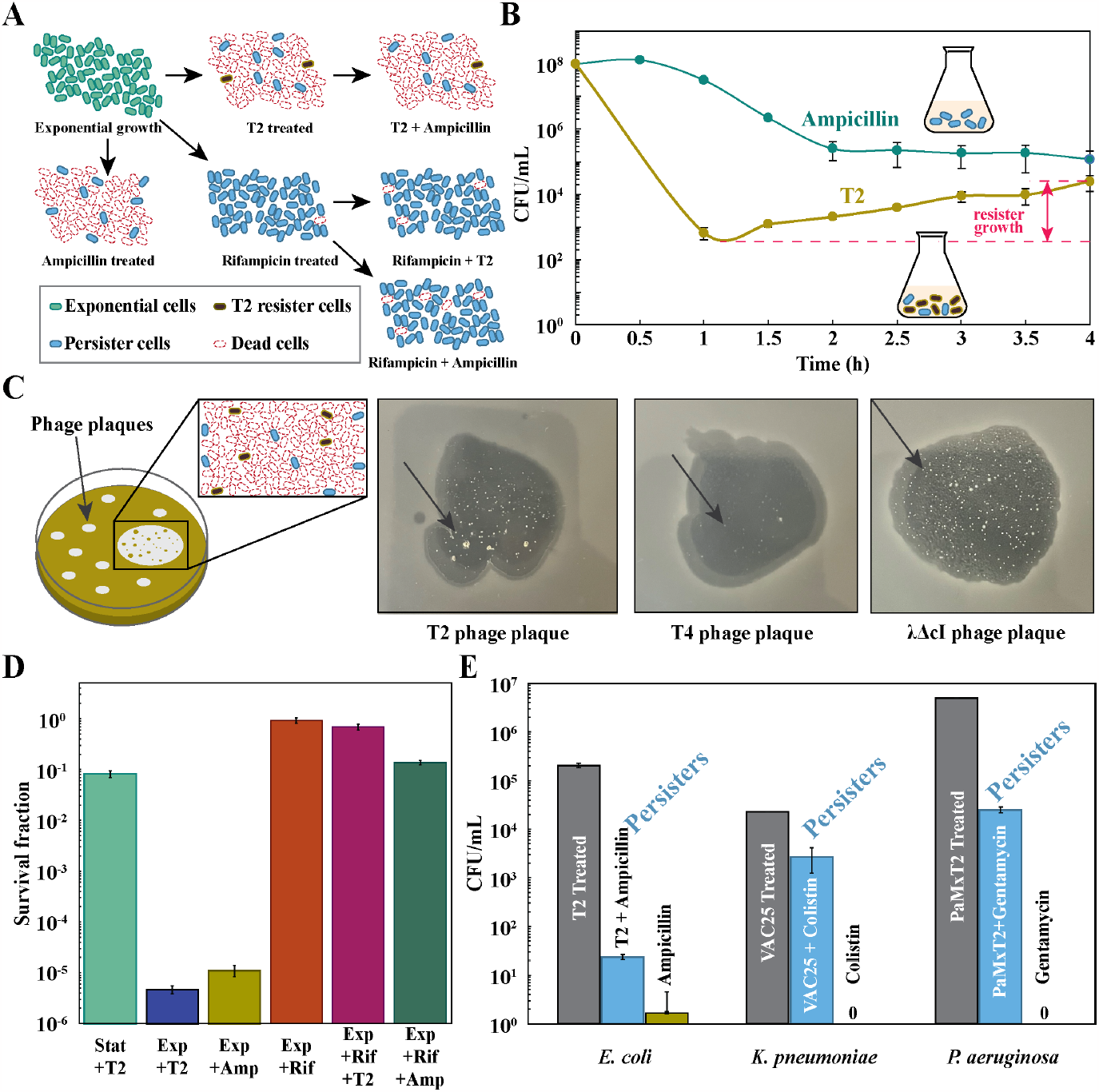
Phages produce persisters. (**A**) Schematic of experiments used to demonstrate that phages produce persister cells. (**B**) Kill curves of exponentially-growing *E. coli* BW25113 treated with ampicillin (100 μg/mL, 10 MIC, blue) or T2 phage (MOI ≈ 0.1, orange). Note the increase in cell density with T2 phage indicates that resistant cells are present along with the persistent cells (with a specific growth rate of 1.2 ± 0.1 h^-1^). (**C**) Double layer TA plates, from left to right, of phages T2, T4, and lambda mutant *cI* infecting *E. coli* BW25113, showing surviving colonies inside the phage inhibition area (indicated with a black arrow). Colonies are visible after one day but allowed to grow for several days for the photo here. (**D**) *E. coli* BW25113 persister cells were formed by rifampicin pre-treatment (30 min, 100 μg/mL), and 0.1 MOI T2 phage was added for 3 h. Exponential cells (turbidity 0.5 at 600 nm) were treated with ampicillin (10 MIC) for 3 h. (**E**) First bar indicates initial cell density after phage attack (10^8^ *K. pneumoniae* cells/mL treated with 0.01 MOI VAC25 phage for one h or 10^8^ *P. aeruginosa* cells/mL treated with 1 MOI PaMx12 phage for one hour), second bar indicates phage and antibiotic treatment (0.01 MOI VAC25 phage for one h followed by 10X MIC of colistin for 3 h for *K. pneumoniae* or 1 MOI PaMx12 phage for one hour followed by 10X MIC of gentamicin for 3 h), and third bar indicates antibiotic treatment alone (10X MIC of colistin for 3 h added to 10^4^ cells/mL for *K. pneumoniae* or 10X MIC of gentamicin for 3 h added to 10^6^ cells/mL for *P. aeruginosa*). One average deviation shown.

### T2-treated cells become persisters

To determine whether the bacteria that survive lytic phage attack are resistant or persistent, we made plaques formed by T2, T4, and lambda *cI* phages and isolated surviving cells from inside the plaques (**Fig. 1c**). The vast majority of cells die, as evidenced by the clearing of lawns when plaques form. We then purified the cells from the phages by streaking twice on LB plates for single colonies. Some purified cells from the T4 plaque were mucoid; hence, mutation clearly occurred with some of the isolates. For both T4 and lambda *cI*, only resistant strains were identified with increases in survivability up to 5 x 10^5^-fold for T4 and 1 x 10^5^-fold for lambda *cI* (**Table 1a**). In contrast, with T2, all four colonies tested remained sensitive to the lytic phage (**Table 1a**). Therefore, the colonies produced in the plaques formed after T2 addition were made by persister cells, since these cells remain sensitive to the phage like the wild-type strain.

**Table 1.**
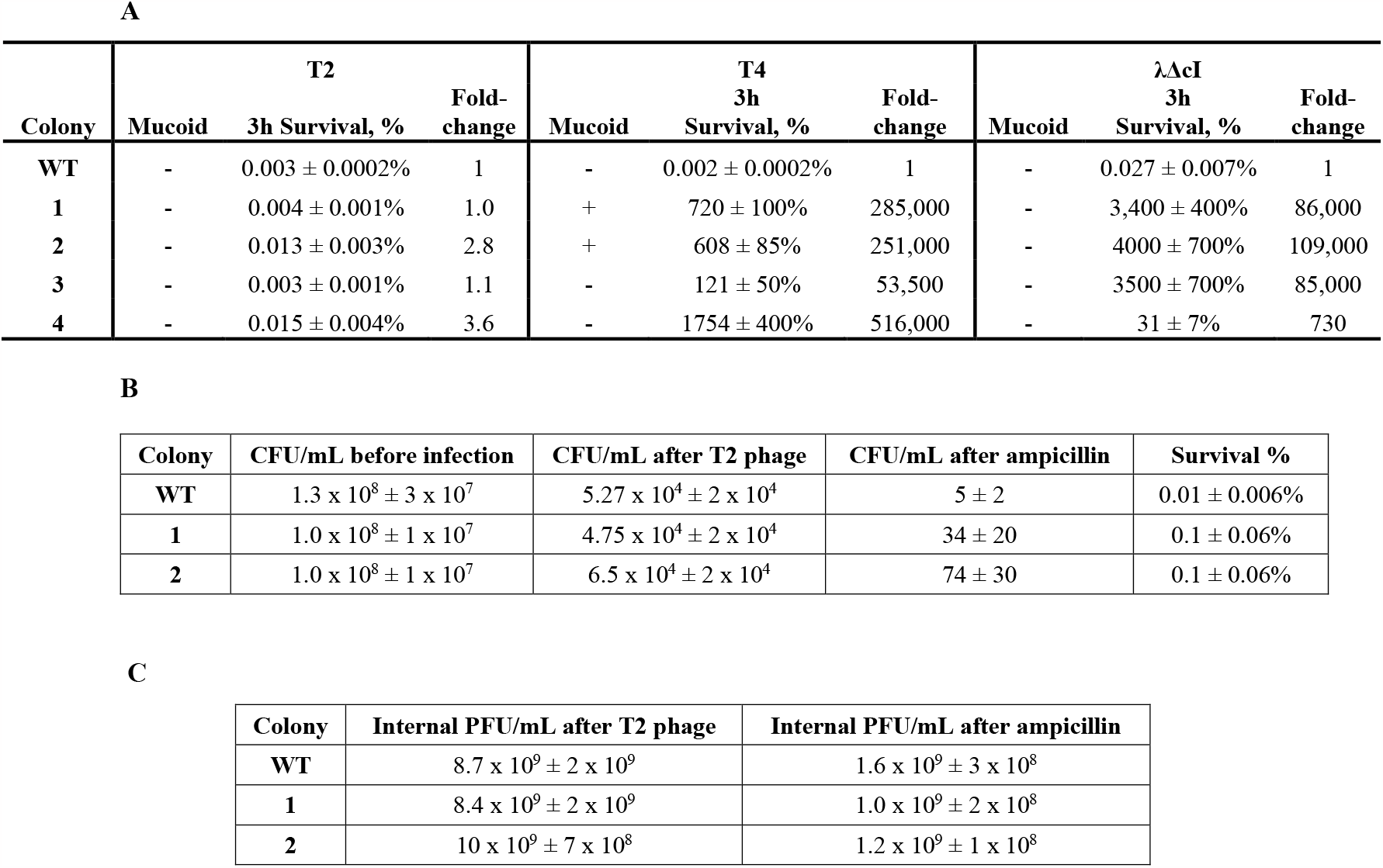
Persister cells of BW25113 against phages. **(A)** Survival of BW25113 against phages, mucoid phenotype of the colonies (+/-), and survival percentage of the colonies isolated from inside the phage plaque when they are retested against the original phages (MOI ∼ 0.1). Survival fold-change is based on that of the wild-type (WT). **(B)** Persister cells of BW25113 (WT), colony 1 and 2 from inside the T2 phage plaque, showing CFU/ml at each step of the persister assay and survival (0.01 MOI+). **(C)** Phages inside cells, measured after each step of the persister assay for BW25113 (WT), colony 1 and 2 from inside the T2 phage plaque.

To enumerate the number of cells in the plaque region that survive T2 infection, we determined the percentage of surviving cells in plaques by averaging the number of cells in nine volumes equivalent to that of the plaques formed by T2 phage in 24 h and found 0.00006 ± 0.00002% survive. For example, 203 colonies formed out of roughly 2.6 x 10^8^ ± 2 x 10^8^ cells in one plaque. Since the cells that survived T2 phage were clearly not resistant; i.e., had no phage resistance (**Table 1a**), we focused on T2 phage for the remaining experiments with *E. coli*.

### Sequencing

To corroborate that the cells that survived phage attack were persistent, we sequenced the whole genome of the parental strain along with four colonies from inside each phage plate for T2, T4, and lambda *cI* using Illumina MiSeq (**Table 2**). Unlike for the resistant T4 and lambda *cI* colonies, which had up to 79 single nucleotide polymorphism mutations (**Table 2**) and which were shown to be resistant (**Table 1a**), there were far fewer mutations in the whole genome of the T2 colonies from within the plaque that were not resistant (as few as 1 coding change, **Table 2**). Moreover, for C1 from a T2 plaque, there is a single coding mutation, in *pinR*; however, this is probably not a gain-of-function mutation as it is as a conservative substitution practically at the N terminus (R3Q so aa position #3 out of 196 aa’s). In addition, deleting *pinR* did not change the growth rate in rich medium but caused a 330-fold increase in T2 sensitivity, so PinR is necessary for host defense against T2. Also, since the colonies in the plaques were sequenced after two steps of 24 h growth in plaques and 16 h of regrowth in liquid media (total 64 h growth), the mutations in the persister cells probably occurred *after* the persistence phase induced by T4 infection.

**Table 2.**
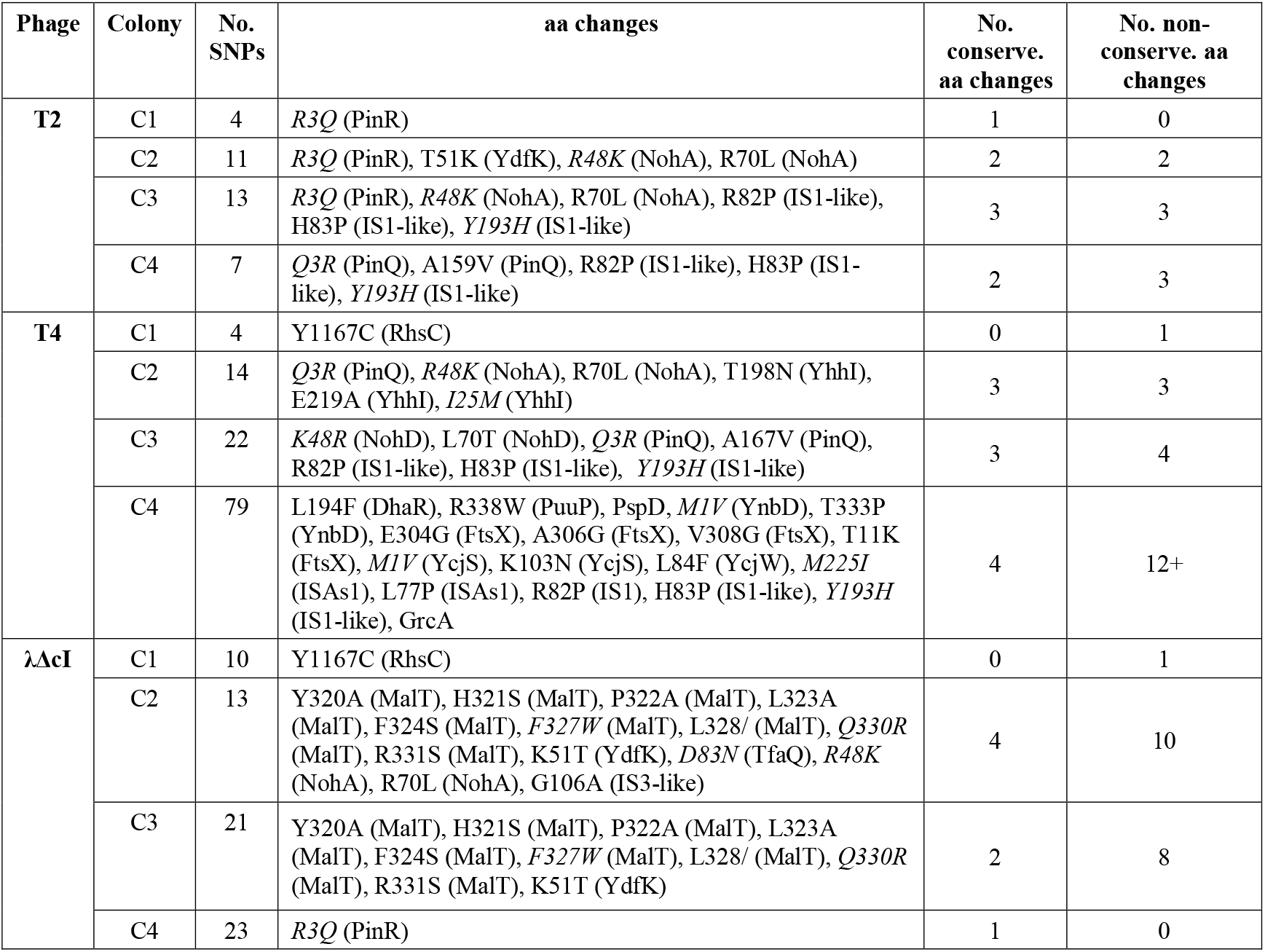
Summary of the SNPS and coding change mutations in the colonies isolated from inside the phage inhibition areas with T2, T4, and λΔcI phages. Coding changes in italics are conservative mutations; i.e., the amino acid is substituted for another in the same group. Those proteins without a specific amino acid change indicate changes throughout the protein. SNPs: single nucleotide polymorphisms, “No.”: number, and “conserve.”: conservative (i.e., similar aa substitutions).

### Cells surviving T2 phage attack are antibiotic persisters

To provide additional evidence that the *E. coli* cells isolated from plaques formed by T2 are persisters, we tested antibiotic sensitivity of the surviving cells after T2 treatment. Critically, treatment of exponentially-growing cells (turbidity of 0.5 at 600 nm) with T2 phage (0.01 MOI) for one hour (to induce persistence) followed by washing to remove external T2 phage and treatment with 10X the minimum inhibitory concentration (MIC) of ampicillin (to lyse non-persister cells), revealed 0.1% of the surviving cells (colonies 1 & 2) and 0.01% of the wild-type (**Table 1b**) become persistent during phage attack; i.e., these cells survived treatment with a lethal ampicillin concentration.

Since we get a small number of persisters after T2 treatment (0.01 to 0.1%), we hypothesized that internal T2 kills some of persister cells upon their resuscitation on the plates used to count the number of surviving cells. To investigate this, we added chloroform to washed cells after T2 treatment and found ∼10^10^ PFU (plaque formation units)/ml from internalized phage for the wild-type and surviving cells isolated from the inside of plaques (**Table 1c)** as well as found 10^9^ PFU (plaque formation units)/ml of internalized phage after subsequent treatment with ampicillin. These results demonstrate clearly the presence of T2 phage inside the cells. Thus, we conclude that some cells probably die upon waking and therefore the number of persister cells is even larger. Corroborating this, anomalous colony shapes (with areas of absent cells in circular colonies) were seen on the plates when quantifying the number of surviving cells.

### Persister cells survive antibiotic treatment

Since T2 infection creates persister cells, we hypothesized that persister cells would withstand T2 infection better than exponentially-growing cells. Hence, we used rifampicin to form *E. coli* persister cells and lysed non-persister cells with ampicillin^10, 11, 13, 24, 25, 29^. This method leads to a 10^5^-fold increase in persister cells and has been vetted nine ways^11, 29^ by us to show the cells generated are *bona fide* persister cells, and it has been used by over 33 groups to induce persistence.

We found a 212,500-fold reduction in cell density when T2 was added to exponentially-growing cells (turbidity ∼ 0.5, MOI of 0.1, **Fig. 1d**). This reduction in cell density was comparable to that of ampicillin treatment to exponentially-growing cells (89,500-fold, **Fig. 1d)**. However, as expected, we found that persister cells were resilient to T2 infection; i.e., there was a 137,000-fold reduction in the ability of T2 to propagate with persister cells compared to exponentially-growing cells treated with T2 (**Fig. 1d**). We also tested the ability of stationary-phase cells (turbidity ∼ 2.5) to withstand T2 infection and found there was an 8-fold increase in infection for stationary-phase cells relative to persister cells (**Fig. 1d**); therefore, persister cells are distinct from slowly-growing stationary cells. Hence, persister cells are dramatically less sensitive to T2 phage infection.

### *K. pneumoniae* and *P. aeruginosa* form persister cells after phage attack that may be eradicated by mitomycin C

We also hypothesized that induction of persistence during lytic infection would be a general phenomenon. To explore whether persister cells are formed after phage attack in non-*E. coli* strains, we investigated whether phages induce persistence in *K. pneumoniae* and *P. aeruginosa*. For *K. pneumoniae*, we found that after treatment of 10^8^ cells/mL with VAC25 phage (0.01 MOI) for one hour, of the remaining viable cells after phage attack (10^4^ cells/mL), 11 ± 4% of the cells were persistent as shown by survival with 10X MIC of colistin for 3 h (**Fig. 1e)**. In comparison, starting with the same initial cell density (10^4^ cells/mL) but omitting phage pretreatment, 0 ± 0% the cells survived 3 h 10X MIC of colistin treatment (**Fig. 1e**). Furthermore, since mitomycin C kills persister cells^5, 12,^ we tested this and found that mitomycin C eradicated the *K. pneumoniae* persister cells that were formed after VAC25 attack.

For *P. aeruginosa*, we found that after treatment of 10^8^ cells/mL with PaMx12 phage (1 MOI) for one hour, of the remaining viable cells after phage attack (10^6^ cells/mL), 0.5 ± 0.14% cells were persistent as shown by survival with 10X MIC of gentamicin for 3 h. In comparison, starting with the same initial cell density (10^6^ cells/mL) but omitting phage pretreatment, 0 ± 0% the cells survived 3 h 10X MIC of gentamicin. In addition, we found that 40% of the cells from the middle of plaques formed in soft agar are persistent, whereas 60% are resistant; i.e., 60% of the cells from the plaques developed mutations that reduced plaque formation in a subsequent assay with phage PaMx12. Hence, both *K. pneumoniae* and *P. aeruginosa* form persister cells upon lytic infection.

For comparison, we performed this set of experiments for *E. coli* (**Fig. 1e**). We also found that the addition of T2 phage produces persister cells; i.e., cells that survive 3 h of lethal ampicillin treatment.

## DISCUSSION

Our results show persisters arise in phage plaques based on five lines of evidence: (i) kill curves show *E. coli* cells surviving phage infection are similar to cells that survive antibiotics, (ii) T2-treated *E. coli* cells isolated from plaques remain sensitive to the phage to the same extent as the wild-type, (iii) sequencing (after 64 h) shows few mutations for these sensitive *E. coli* cells isolated from plaques (compared to resistant cells from T4 and lambda *cI*), (iv) T2 addition to *E. coli* persister cells results in dramatically less killing compared to exponentially-growing cells, and (v) cells that survive T2 attack become tolerant to lethal antibiotic concentrations (10 MIC). Corroborating the *E. coli* data, we found both *K. pneumoniae* and *P. aeruginosa* formed persister cells after phage attack; hence, induction of persistence during lytic infection is a general phenotype. **Fig. 2** summarizes our results.

**Fig. 2.**
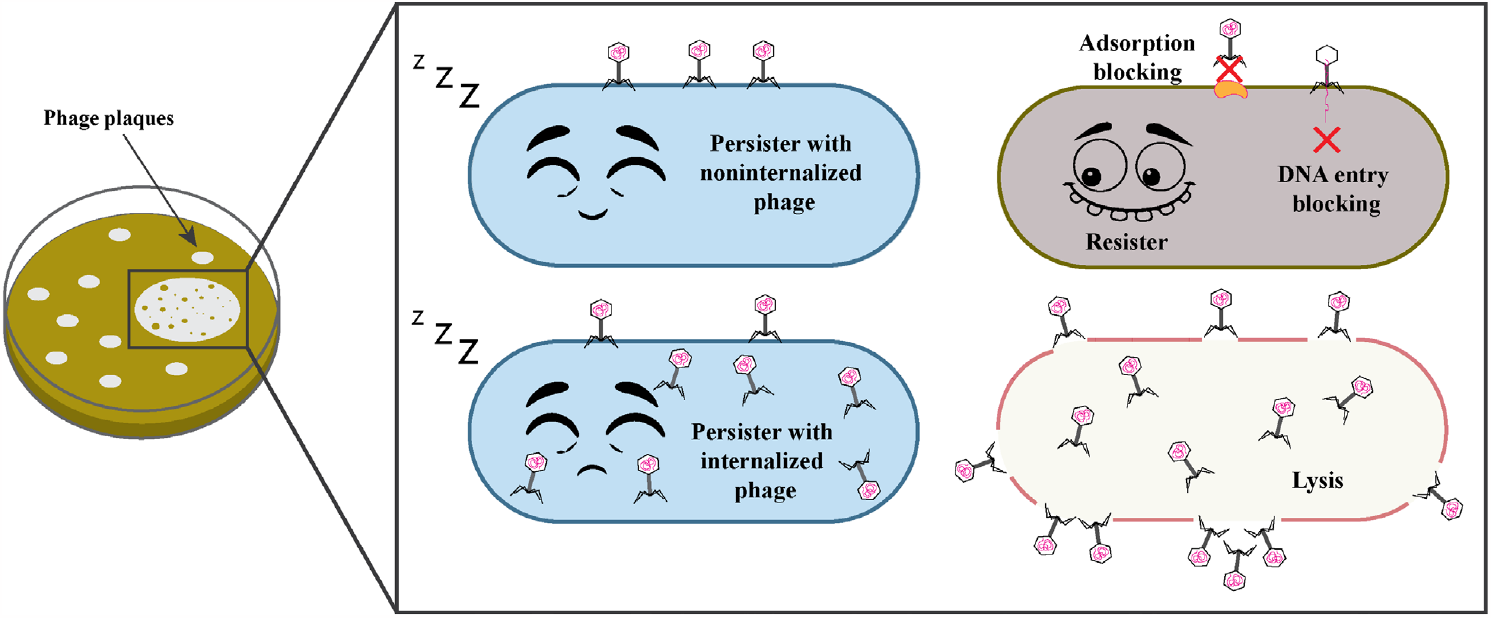
Schematic of model for cells in plaques.

The genesis of the persister cells that form during phage infection is likely a result of increased guanosine tetraphosphate that leads to ribosome dimerization^24, 25, 29^. Moreover, the increase in guanosine tetraphosphate may be an indirect result of phage defense mechanisms like toxin/antitoxin and restriction/modification systems.

Our results with virulent lambda agree well with previous results where it was shown phages with a *cI60* mutation kill persister cells^20^ since we did not isolate readily persister cells from plaques formed with virulent lambda phage. However, unlike previous results with a virulent, superinfecting lambda phage where no survivors were detected^20^, our results are in stark contrast, as we observed survival and resuscitation of persister cells after superinfection with T2 phage both from plaques, from suspension cultures, and pre-formed persister cells. Similar survival of a small population of *E. coli* cells was reported previously with T4 phage although it was concluded these ‘hibernators’ wake and are lysed by T4 phage, and the hibernators were not tested for persistence^4^. Similarly, a ‘phage tolerance response’ is elicited in non-infected *Bacillus subtilis* by an unknown product of cell lysis, but this response is not persistence and is, instead, is a SigX-mediated stress response ^27^.

In prior work, it was suggested that phages could be used to target persister cells^20^, whereas we conclude it may be better to combine phages (which will generate persister cells) with anti-persister compounds like mitomycin C, which kills persister cells by cross-linking their DNA^12^. Mitomycin C has been shown to kill numerous pathogenic persisters^5, 12^ and has been shown to work well when combined with phage therapy for *Klebsiella* spp.^19^.

Just like resistance to phage, persistence should be considered as a possible deleterious outcome of phage therapy. In addition, our results show that phage infection is similar to other stresses (e.g., antibiotics^13^, oxidative stress^9^, and starvation^10^) that causes persistence.

## MATERIALS AND METHODS

### Bacteria and growth conditions

Bacteria and phages are shown in **Table S1**, and cells were cultured at 37oC in lytic broth^15^ (LB). The BW25113 strain was checked to ensure that it was not a lambda lysogen by PCR using *tomB* primers as a positive control (forward 5’-CGATTACCTGACTTCCGCCA and reverse 5’-TCATGGCTGGGTAAACGACC) and the *cI* lambda primers to check for the presence of lambda (forward 5’-CACCCCCAAGTCTGGCTATG, and reverse-5’ACCAAAGGTGATGCGGAGAG).

### Kill curves

The kill curve assays to verify *E. coli* persister cells were formed during ampicillin and T2 phage treatment was performed by treating late exponential cells (turbidity at 600 nm of ∼ 0.5) with 100 μg/ml ampicillin for 4 hours with shaking at 250 rpm in a 15 mL flask or by treating with T2 phage at a MOI of 0.1 for 4 h. Samples were taken every 30 min, washed with phosphate-buffered saline (PBS, 8 g NaC1, 0.2 g KC1, 1.15 g Na_2_HPO_4_, and 0.2 g KH_2_PO_4_ in dH_2_O 1000 mL), and 100 μL was serially diluted with PBS to determine the number of viable cells.

### Sensitivity assay for plaque-derived bacteria and sequencing mutant

*E. coli* strains were streaked on fresh LB agar plates and incubated overnight. A single colony was inoculated into 15 mL of LB broth and incubated with shaking (250 rpm) for 16 h. The overnight culture (100 μL) was used to create a double layer phage plaque with a 20 μL drop of T2, T4 or lambda (cI mutant) phage and incubated overnight. The next day, colonies from inside the inhibition-lysis area of the phage plaque were streaked twice on fresh LB plates to purify strains and to remove phage. Single colonies were cultured overnight in LB, then diluted 100X and grown to a turbidity of 0.5 at 600 nm. T2, T4 or lambda (cI mutant) phage was added at a MOI ∼ 0.1 for three hours, then the cells were washed twice with PBS and enumerated on LB plates to measure survival in the presence of phage. Similarly, for the *pinR* mutant, T2 phage was added (0.1 MOI) and the number of viable cells was determined after 3 h.

To estimate the number of insensitive bacteria in plaques, T2 phage was added at ∼0.1 MOI via a 20 μL drop to double-layer plates, then the number of surviving cells in the plaque was counted after overnight incubation. For nine areas approximately the size of the plaques, the number of cells was determined by resuspending in 1 mL of PBS and using the drop assay.

### Persister assays

*E. coli* persister cells were generated by 30 min rifampicin pretreatment (100 μg/mL) followed by ampicillin treatment (10 MIC) to lyse non-persister cells as described previously^10, 11, 13, 24, 25, 29^. After rifampicin treatment and washing twice with PBS, T2 phage was added for 3 h (0.1 MOI), and cell viability was tested by washing twice with PBS to remove external phage and enumerating bacteria via the drop assay.

To test for the induction of persistence during T2 treatment (but without rifampicin pretreatment), survival in antibiotic was tested, by using single *E. coli* colonies that were cultured overnight in LB, then diluted 100X and grown to a turbidity of 0.5 at 600 nm. T2 phage was added (MOI ∼ 0.01), cells were incubated for an hour, then 5 mL of cells were harvested by centrifugation at 5,000 rpm for 10 min, washed with PBS, and resuspended in 5 mL of LB containing 100 μg/mL of ampicillin (10 MIC). The cultures were incubated for 3 h with shaking at 250 rpm in a 15 mL culture tube, washed twice with PBS, then 100 μL were serially diluted with PBS to determine the number of viable cells. For enumerating internal phage during the persister assay, the cell samples were washed twice with PBS to remove external phage and treated with 1% chloroform, then serially diluted with phage buffer.

### *Klebsiella pneumoniae* and *Pseudomonas aeruginosa* persister assays after phage attack

*K. pneumoniae* strain obtain from blood infection^18^ was infected with DNA lytic phage vB_KpnP-VAC25 ^3^ at MOI 0.01, washed with PBS, and contacted with 10 MIC colistin (10 μg/mL) or 5 MIC mitomycin C (10 μg/mL). *P. aeruginosa* PAO1 from the Gloria Soberón collection at UNAM was infected with phage PaMx12 (DNA lytic phage, sequence https://www.ncbi.nlm.nih.gov/nuccore/386649691) at MOI 1 and surviving colonies in the center of the plaques were isolated after 20 h and tested for phage sensitivity to determine if the cells were resistant or persistent to phage. To confirm persistence, cells were contacted with phage followed by lethal (10 MIC, gentamycin at 10 μg/mL for *P. aeruginosa*).

### Sequencing

Genomic *E. coli* DNA was purified using the Qiagen DNA isolation kit following the manufacturer instructions. The quality of samples was quantified by nanodrop and Qubit. Samples were sequenced using Illumina MiSeq by the Genomics Core Facility at the Pennsylvania State University. The sequences were assembled and analyzed using bv-brc.org^17^.

## ACKNOWLEDGEMENTS

This work was supported by both a Fulbright Scholar Fellowship and a Xunta de Galicia Postdoctoral Grant for LFG. This study has been funded by the Instituto de Salud Carlos III (ISCIII) through projects PI19/00878 and PI22/00323 and co-funded by the European Union, by a Personalized and Precision Medicine Grant from the Instituto de Salud Carlos III (MePRAM Project, PMP22/00092) and by the Study Group on Mechanisms of Action and Resistance to Antimicrobials, GEMARA (SEIMC). R.G.-C. was supported by DGAPA, PAPIIT-UNAM (grant number IN200121). D.H.-M. gratefully acknowledges the Programa de Maestría y Doctorado en Ciencias Bioquímicas, UNAM and the MSc scholarship 09970 from Consejo Nacional de Ciencia y Tecnología (CONACyT, México). We appreciate the assistance of Erin Essington on editing the figures.

## Supporting Information

**Table S1.**
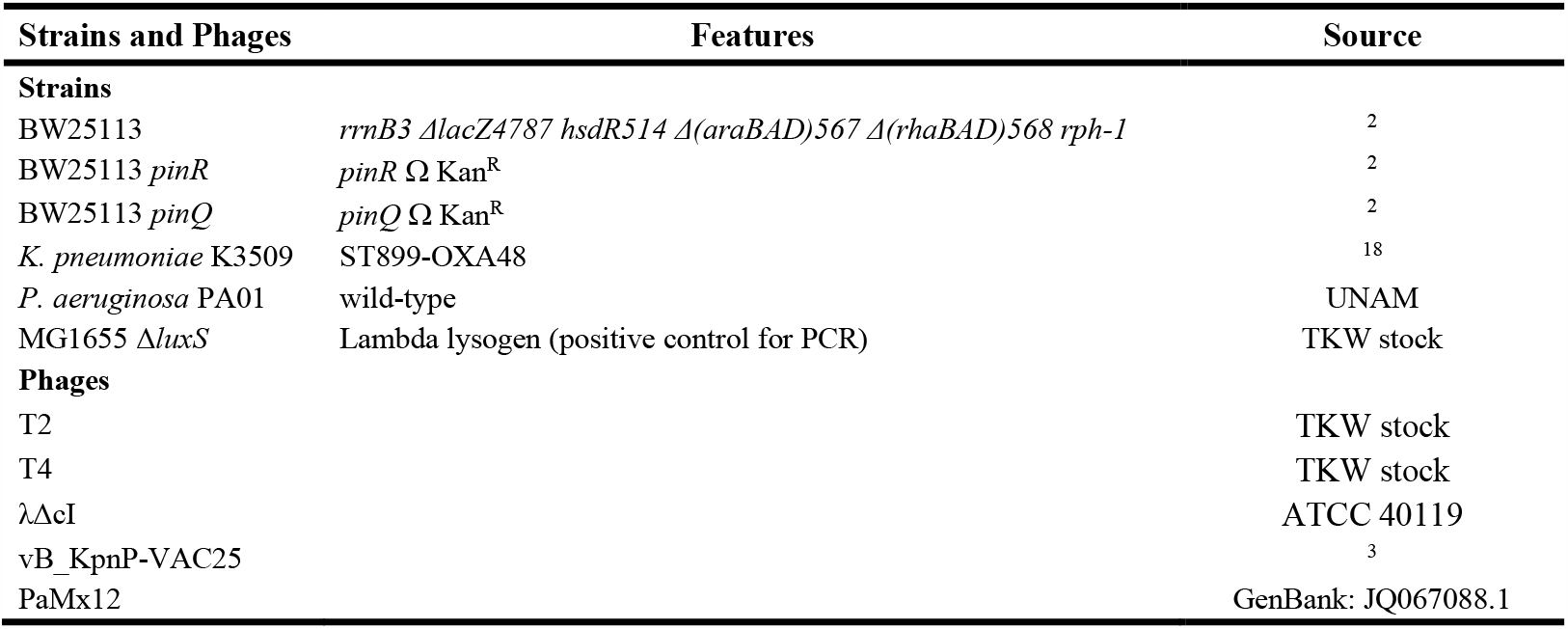
*E. coli* bacterial strains and plasmids utilized. Cm^R^ is chloramphenicol resistance, and Kan^R^ is kanamycin resistance.

## REFERENCES

1. Abedon ST. 2018. Detection of Bacteriophages: Phage Plaques, p. 1–32. In Harper DR, Abedon ST, Burrowes BH, McConville ML (ed.), Bacteriophages: Biology, Technology, Therapy. Springer International Publishing, Cham.

2. Baba T, Ara T, Hasegawa M, Takai Y, Okumura Y, et al. 2006. Construction of Escherichia coli K-12 in-frame, single-gene knockout mutants: the Keio collection. Mol Sys Biol 2:2006.0008.

3. Bleriot I, Blasco L, Pacios O, Fernández-García L, López M, et al. 2023. Proteomic Study of the Interactions between Phages and the Bacterial Host Klebsiella pneumoniae. Microbiol Spectr 11:e0397422.

4. Bryan D, El-Shibiny A, Hobbs Z, Porter J, Kutter EM. 2016. Bacteriophage T4 Infection of Stationary Phase E. coli: Life after Log from a Phage Perspective. Front Microbiol 7: 1391.

5. Cruz-Muñiz MY, López-Jacome LE, Hernández-Durán M, Franco-Cendejas R, Licona-Limón P, et al. 2017. Repurposing the anticancer drug mitomycin C for the treatment of persistent Acinetobacter baumannii infections. Int J Antimicrob Agents 49:88–92.

6. Fernandez-Garcia L, Song S, Kirigo J, Petersen ME, Tomas M, et al. 2023. Toxin/Antitoxin Systems Induce Persistence and Work in Concert with Restriction/Modification Systems to Inhibit Phage. bioRxiv:2023.2002.2025.529695.

7. Fernández-García L, Tomás M, Wood TK. 2023. Ribosome inactivation by Escherichia coli GTPase RsgA inhibits T4 phage. Frontiers in Microbiology 14: 1242163.

8. Fernández-García L, Wood TK. 2023. Phage-Defense Systems Are Unlikely to Cause Cell Suicide. Viruses 15:1795.

9. Hong SH, Wang X, O’Connor HF, Benedik MJ, Wood TK. 2012. Bacterial persistence increases as environmental fitness decreases. Microb Biotechnol 5:509–522.

10. Kim J-S, Chowdhury N, Yamasaki R, Wood TK. 2018. Viable But Non-Culturable and Persistence Describe the Same Bacterial Stress State. Environ Microbiol 20:2038–2048.

11. Kim J-S, Yamasaki R, Song S, Zhang W, Wood TK. 2018. Single Cell Observations Show Persister Cells Wake Based on Ribosome Content. Environ Microbiol 20:2085–2098.

12. Kwan BW, Chowdhury N, Wood TK. 2015. Combatting Bacterial Infections by Killing Persister Cells with Mitomycin C. Environ Microbiol 17 4406–4414.

13. Kwan BW, Valenta JA, Benedik MJ, Wood TK. 2013. Arrested protein synthesis increases persister-like cell formation. Antimicrob Agents Chemother 57:1468–1473.

14. Little JS, Dedrick RM, Freeman KG, Cristinziano M, Smith BE, et al. 2022. Bacteriophage treatment of disseminated cutaneous Mycobacterium chelonae infection. Nature Comm 13:2313.

15. Maniatis T, Fritsch EF, Sambrook J. 1982. Molecular Cloning: A Laboratory Manual. Cold Spring Harbor, Cold Spring Harbor.

16. Murray CJL, Ikuta KS, Sharara F, Swetschinski L, Robles Aguilar G, et al. 2022. Global burden of bacterial antimicrobial resistance in 2019: a systematic analysis. The Lancet 399:629–655.

17. Olson RD, Assaf R, Brettin T, Conrad N, Cucinell C, et al. 2023. Introducing the Bacterial and Viral Bioinformatics Resource Center (BV-BRC): a resource combining PATRIC, IRD and ViPR. Nucleic Acids Res 51:D678–d689.

18. Pacios O, Fernández-García L, Bleriot I, Blasco L, Ambroa A, et al. 2021. Phenotypic and Genomic Comparison of Klebsiella pneumoniae Lytic Phages: vB_KpnM-VAC66 and vB_KpnM-VAC13. Viruses 14: 6.

19. Pacios O, Fernández-García L, Bleriot I, Blasco L, González-Bardanca M, et al. 2021. Enhanced Antibacterial Activity of Repurposed Mitomycin C and Imipenem in Combination with the Lytic Phage vB_KpnM-VAC13 against Clinical Isolates of Klebsiella pneumoniae. Antimicrob Agents Chemother 65:e00900–00921.

20. Pearl S, Gabay C, Kishony R, Oppenheim A, Balaban NQ. 2008. Nongenetic Individuality in the Host–Phage Interaction. PLOS Biology 6:e120.

21. Perlemoine P, Marcoux PR, Picard E, Hadji E, Zelsmann M, et al. 2021. Phage susceptibility testing and infectious titer determination through wide-field lensless monitoring of phage plaque growth. PLOS ONE 16:e0248917.

22. Postic B, Finland M. 1961. Observations on Bacteriophage Typing of Pseudomonas aeruginosa. The Journal of Clinical Invest 40:2064–2075.

23. Pouillot F, Chomton M, Blois H, Courroux C, Noelig J, et al. 2012. Efficacy of Bacteriophage Therapy in Experimental Sepsis and Meningitis Caused by a Clone O25b:H4-ST131 Escherichia coli Strain Producing CTX-M-15. Antimicrob Agents Chemother 56:3568–3575.

24. Song S, Wood TK. 2020. Persister Cells Resuscitate via Ribosome Modification by 23S rRNA Pseudouridine Synthase RluD. Environ Microbiol 22:850–857.

25. Song S, Wood TK. 2020. ppGpp Ribosome Dimerization Model for Bacterial Persister Formation and Resuscitation. Biochem Biophys Res Com 523:281–286.

26. Strathdee SA, Hatfull GF, Mutalik VK, Schooley RT. 2023. Phage therapy: From biological mechanisms to future directions. Cell 186:17–31.

27. Tzipilevich E, Pollak-Fiyaksel O, Shraiteh B, Ben-Yehuda S. 2022. Bacteria elicit a phage tolerance response subsequent to infection of their neighbors. EMBO J 41:e109247.

28. Williams MC, Reker AE, Margolis SR, Liao J, Wiedmann M, et al. 2023. Restriction endonuclease cleavage of phage DNA enables resuscitation from Cas13-induced bacterial dormancy. Nat Microbiol 8:400–409.

29. Yamasaki R, Song S, Benedik MJ, Wood TK. 2020. Persister Cells Resuscitate Using Membrane Sensors that Activate Chemotaxis, Lower cAMP Levels, and Revive Ribosomes. iScience 23:100792.

